# Artificial contractile actomyosin gels recreate the curved and wrinkling shapes of cells and tissues

**DOI:** 10.1101/2023.03.21.533327

**Authors:** Gefen Livne, Shachar Gat, Shahaf Armon, Anne Bernheim-Groswasser

**Affiliations:** Department of Chemical Engineering, Ilse Kats Institute for Nanoscale Science and Technology, Ben Gurion University of the Negev, Beer-Sheva 84105, Israel; Department of Physics, Weizmann Institute of Science, Rehovot 76100, Israel

## Abstract

Living systems adopt a diversity of curved and highly dynamic shapes. These diverse morphologies appear on many length-scales, from cells to tissues and organismal scales. The common driving force for these dynamic shape changes are contractile stresses generated by myosin motors in the cell cytoskeleton, an intrinsically active filamentous material, while converting chemical energy into mechanical work. A good understanding of how contractile stresses in the cytoskeleton arise into different 3D shapes and what are the selection rules that determine their final configurations still lacks. Aiming to identify the selection rules governing the shapes formed by contractile forces in living systems, we recreate the actomyosin cytoskeleton in-vitro, with precisely controlled composition and initial geometry. A set of actomyosin gel discs, intrinsically identical but of variable initial geometry, spontaneously self-organize into a family of 3D shapes. This process occurs through robust distinct dynamical pathways, without specific pre-programming and additional regulation. Shape selection is encoded in the initial disc radius to thickness aspect ratio, and thus scale-free. This may indicate a universal process of shape selection, that works across scales, from cells to tissues and organelles. Finally, our results suggest that, while the dynamical pathways may depend on the detailed interactions of the different microscopic components within the gel, the final selected shapes obey the general theory of elastic deformations of thin sheets. Altogether, these results provide novel insights on the mechanically induced spontaneous shape transitions in active contractile matter and uncover new mechanisms that drive shape selections in living systems across scales.

**Significance statement:** Living systems adopt a diversity of curved and highly dynamic shapes. These diverse morphologies appear on many length-scales, from cells to organismal scales, and are commonly driven by contractile stresses generated by myosin motors in the cell cytoskeleton. By recreating the actomyosin cytoskeleton in-vitro, with precisely controlled composition and initial geometry, we identify the shape selection rules that determine the final adopted configuration. Specifically, we find that shape selection is scale-free, which may indicate a universal process of shape selection, that works across scales, from cells to tissues and organelles. Altogether, our results provide novel insights on the mechanically induced spontaneous shape transitions in contractile active matter and uncover new mechanisms that drive shape selections in living systems.

## Introduction

Living systems can adopt a variety of curved shapes. These diverse morphologies appear on many length-scales, from cells to organismal scales, underlying processes like cell migration and tissue morphogenesis (1–4). The common driving force for these dynamic shape changes are contractile stresses generated by myosin motors, generated by converting chemical energy into mechanical work. These stresses propagate through the cell cytoskeleton, an adaptive, intrinsically active (5–7), poroelastic contractile filamentous actin network (8, 9). Shape deformations may be guided by topological defects in aligned actin filaments (10) and in arrangements of the active stress-generating components (11).

Due to the complexity of cells, however, the mechanism by which these intrinsically active stresses result in complex 3D shapes during various biologically relevant processes remains poorly understood. A powerful alternative approach is to recreate the actomyosin cytoskeleton outside the cell, using the same actin and myosin building blocks of the cellular cytoskeleton, with precisely controlled concentration of the microscopic constituents and level of activity, to form intrinsically active, crosslinked actomyosin networks. Such networks exhibit spontaneous contraction and self-organization arising in different 2D contraction patterns (12–17) and in the spontaneous wrinkling of thin poroelastic actomyosin gel sheets (18). Thin elastic sheets undergo buckling in response to non-uniform strains, and this phenomenon is harnessed in diverse material science applications to robustly generate complex 3D shapes (19), which can also exhibit oscillatory shape transformation capabilities (20). However, in contrast to uniformly externally stimulated gel sheets (21–24), the contraction and wrinkling instability of actomyosin gel sheets (18) occurs spontaneously without requiring pre-imposed gradients in the network material properties. Despite these advances, a good understanding of how the contractile stresses induced by myosin motors in these networks arise into different 3D shapes and what are the selection rules that determine their final configurations is still lacking.

Here, we identify these selection rules using a set of elastic contractile, intrinsically identical, actomyosin gel discs of variable dimensions. In intent to facilitate the quantification of the changes in the disc radius and thickness, and shape deformation, while resolving the network internal structure, we use relatively large-scale systems. These active gel discs spontaneously self-organize into a family of 3D shapes, notably domes and wrinkled shapes, in response to system initial geometry. This process occurs through robust distinct dynamic pathways, without the need for specific pre-programming and additional regulation. While azimuthal contractility can explain the buckling into a dome, radial contractility can explain the wrinkling deformation. We show that shape selection is encoded in the initial disc radius to thickness aspect ratio, indicating the emergence of shaping scalability. Namely, those systems having the same initial aspect ratio will adopt the same steady state 3D configurations, independent of their actual initial dimensions. This may be indicative of a universal, scale-free process of shape selection. Our results are therefore also relevant to explain both the shape of cells and multi-cellular contractile tissues. Finally, our results suggest that while being intrinsically active systems, the final selected shapes seem to conform to the general theory of elastic deformations of thin sheets.

## Results

Active contractile (poro)-elastic gel discs are prepared as described in(18, 25). In short, a drop of solution containing actin monomers, myosin motors, the strong cross-linker fascin, and ATP is squeezed between two, PEG-passivated, glass coverslips. The spacing ℎ between the two glasses sets the network initial thickness. The system (initial) aspect ratio is defined as 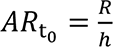, where *R* is the drop radius (Fig. 1A). Network formation is initiated by actin filaments polymerization, which get crosslinked into bundles by fascin. The myosin motors, assembled in aggregates, then bind to the actin bundles to form an interconnected network which locally deform and coarsen with time. Network contraction initiates from the boundary and propagates inwards (18). Buckling follows this initial network forming period, until a mechanically stable buckled state is reached at *t*_end_ (Fig. 1A). The final thickness and (projected) radius are denoted by *b* and *r* (Fig. 1B). Two main types of 3D shape are spontaneously generated: (i) wrinkled shapes of *single* wavy mode with *m* ≥ 2 number of nodes, where *m* = 2 corresponds to a traditional 2-peak saddle, and (ii) large-scale buckled symmetric configurations with a dome shape (*m* = 1).

**Figure 1.**
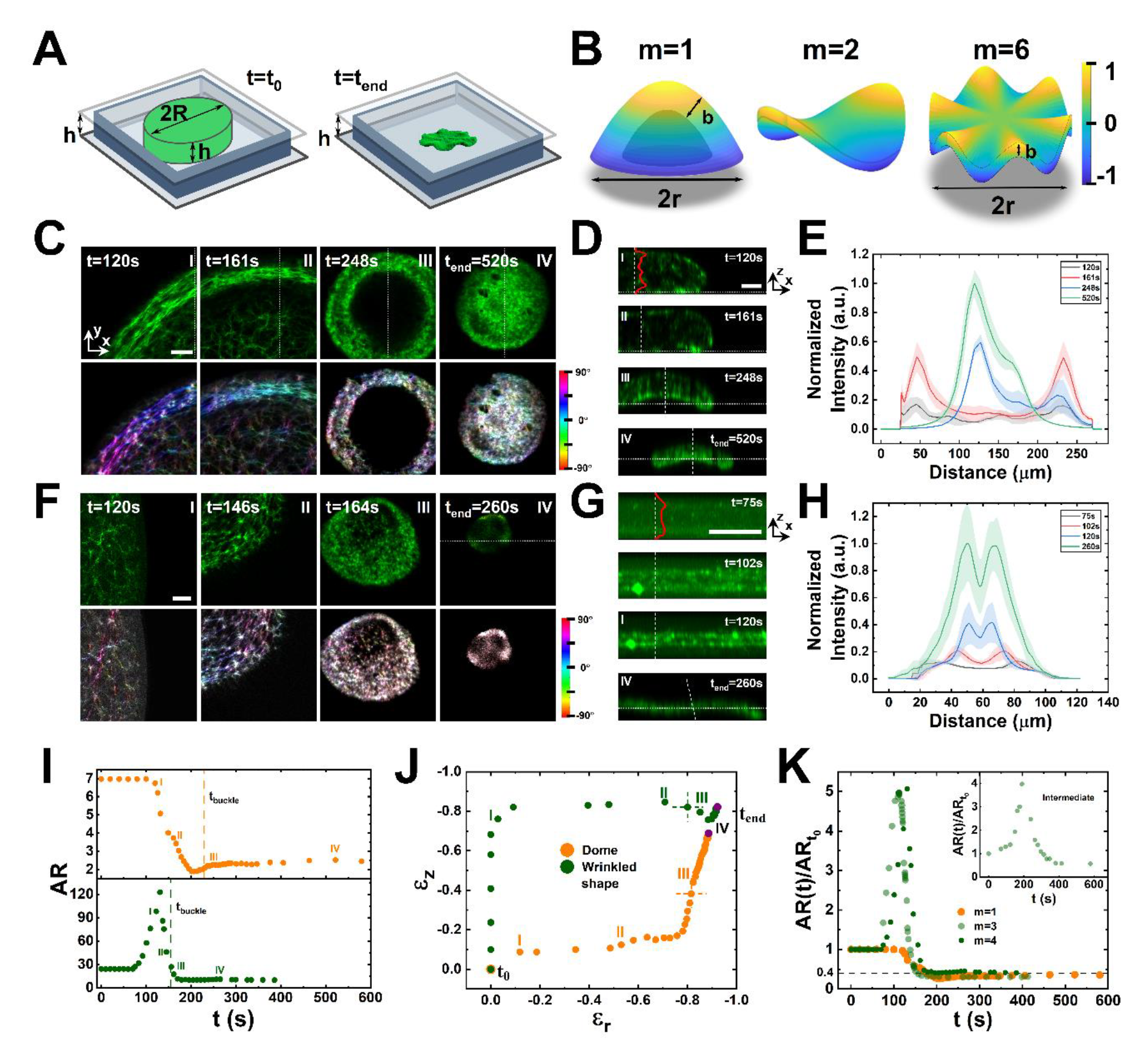
Domes and wrinkled shapes form through distinct dynamic pathways. **A)** (Left) Initial geometry of the actomyosin solution consists of a disc of radius *R* and thickness ℎ. (Right) Final state - shown is a wrinkled shape. **B)** Schematic shapes forming in this study. (Left to right) dome, traditional 2- peak saddle, and a wrinkled shape. *m* – number of nodes, final gel thickness *b* and (projected) radius *r*. Color code: surface high relative to the mid-surface. **(C-E, I-K) Domes shapes. C)** (Upper row) top-view confocal images, measured along the dotted lines in (D) and (bottom row) local filaments orientation, color coded to the orientational angle. **D)** Confocal side-view images measured along the dotted lines in (C). **(E)** Normalized actin density profiles measured normal to the thickness from bottom-to-top, along the dashed lines in (D). The bold lines and surrounding colored area correspond to mean ± SD (*N* = 25 profiles). **(F-H, I-K) Wrinkled shapes.** F-H are as for the dome in C-E. **(G)** Maximal Intensity Projection (MIP) images are provided (*t* ≤ 120 sec). Scale bars: 100 μm. **I)** Aspect ratio, 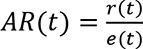, *vs.* time (*r*(*t*) and *e*(*t*) - temporal projected radius and thickness) and **(J)** lateral and vertical strains, 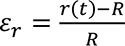 and 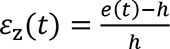 from *t*_0_ to *t*_end_ (purple dots), for the gels in (C,D) and (F,G). The dashed lines and crosses in **(I)** and **(J)** mark the wrinkling (green, *t*_buckle_ = 165 sec) and dome buckling (orange, *t*_buckle_ = 230 sec) onsets; I-IV are as in (C,D) and (F,G), respectively. **(K)** Normalized aspect ratio *AR*(*r*)/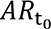 *vs.* time: dome (orange, 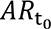 = 7), intermediate (inset, 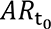 = 8.5), and wrinkled shapes with 3 and 4 nodes (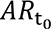 = 23.6 and 26). Steady state value (dashed line) equals 0.4.

### Domes and wrinkled shapes form through distinct dynamic pathways

Dome and wrinkled shapes form through distinct contraction and buckling processes (Fig. 1). The temporal projected radius *r*(*t*) and thickness *e*(*t*) are extracted from confocal microscopy images (Figs. S1-S4), from which the variation in gel aspect ratio *AR* (*t*) = *r*(*t*)/*e*(*t*) and radial and vertical strains, 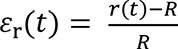 and 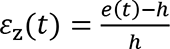 are derived (see Methods and SI).

For all systems, the dynamics are characterized by two main contraction phases. For systems forming dome shapes (Fig. 1C-E), in-plane radial contraction is initially more significant than vertical contraction (Figs. S1, S2, Movie S1), while the second phase is governed by vertical contraction, with only minor changes in projected radius. The gradual decrease in aspect ratio with time (Fig. 1I and 1J, orange dots) and the radial *vs.* vertical strains profile (Fig. 1J), orange dots) are both reflecting these changes in thickness and projected radius with time.

The global changes in gel dimensions are accompanied by the local enrichment of the actin density at the gel periphery, notably, at the top and bottom surfaces (Fig. 1E). This local enrichment may be attributed to network poroelasticity and to the outward water flow generated by gel contraction(18). Also, a *pronounced* azimuthal alignment of the actin bundles at the gel periphery accompanies radial contraction, with the bulk gel remaining isotropic (Fig. 1C, bottom row and Movie S2). This annular orientational order persists as contraction advances, though is more difficult to resolve as the network densifies. As radial contraction advances the system spontaneously buckles into a dome shape, through which the disc center adopts a non-vanishing curvature (*t*_buckle_), which breaks the system up/down symmetry. The buckling instability is preceded by a local deformation of the gel close to the disc edge, which propagates inwards concomitant to radial contraction, turning the local deformation into a global one (Figs. 1D, S1B, Movie S1). The disc buckling also turns the initial symmetric distribution of the actin density across the thickness into an asymmetric one (Fig. 1E), such that the top and bottom surfaces are no longer equivalent. After buckling onset, the gel continues to contract primally along its thickness at slow pace, until a mechanical stable state is reached (*t*_end_). Among the various dome shapes analyzed, 60% point upwards (*N* = 10) suggesting that the top/down symmetry is spontaneously broken.

For systems evolving into wrinkled shapes, gel contraction initiates in vertical direction through which a thin sheet, enriched in actin density at the top and bottom surfaces, is formed (Figs. 1G, 1H, S3A, Movie S3). The thin sheets then contract radially (Fig. 1F). Despite the overall 10-fold decrease in sheet radius, which promotes a corresponding increase in gel density, the thickness remains unchanged, except for small fluctuations (10%) due to recoiling effects (Fig. S4). As radial contraction advances wrinkles spontaneously develop at the sheet periphery (*t*_buckle_), after which contraction proceeds slowly until a mechanically stable state is reached (*t*_end_) (Figs. 1F, 1G, S4). The actin density remains symmetrically distributed across the thickness throughout radial contraction and wrinkling phases, with the sheet top and bottom, surfaces being equivalent (Fig. 1H), which suggests that in-plane gradients develop in the sheet and drives its wrinkling. While azimuthal ring contractility can explain the buckling into a dome, as buckling due to such contraction follows the largest wavelength mode, as in any elastic object, radial contractility can explain the wrinkling deformation (11). The gel sheet remains structurally isotropic, except for a vanishingly small, aligned region of the actin bundles at the sheet periphery (Figs. 1F - bottom row, S3B, Movie S4). This alignment combined with the tension-dependent catch-bond of myosin (26), may suffice for such in-plane radial gradients to develop in the sheet.

The observed non-monotonous variation in aspect ratio reflects the temporal changes in the measured thickness and projected radius (Figs. 1I, green dots). It also explains the inversed radial *vs.* vertical strains profile, which exhibits a much-pronounced time separation between the vertical and radial, dominated contraction phases compared to domes (Fig. 1J, green dots). *All* systems forming wrinkled shapes exhibit similar contraction patterns (Fig. 1K), which interestingly, also holds for, the very few, systems forming intermediate shapes (Fig. 1K, inset). Even more surprisingly, all systems, independently of their final configuration, exhibit a similar net reduction in their aspect ratio of 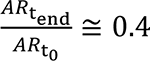 (Fig. 1K).

### Gaussian curvature distribution of the curved 3D configurations

The geometry of the domes and wrinkled shapes can be characterized by calculating the local Gaussian curvature, *K*(*x*, *y*), an intrinsic property of non-flat surfaces which also provides a measure of the local in-plane compression/stretching through Gauss theorem (27). To extract the Gaussian curvature, the topography *z*(*x*, *y*) of the gels’ upper surface is constructed from a stack of *xy* cross-section images, then fitted to a 2D polynomial function (Fig. 2A and SI). This smoothed surface is assumed to reliably represent the shape and local curvatures of the mid-surface. This is explicitly true for wrinkled shapes which form thin sheets (i.e., *b*/*r* ≪ 1). For domes, which are relatively thicker (i.e., *b*/*r* < 8), these values provide a lower bound estimate of the local and average, Gaussian curvatures. The local Gaussian curvature is computed at each point along this surface (Fig. 2B) from which its averaged value, 〈*K*〉, is calculated (Fig. 2C, dashed line). Plotting the distribution of the Gaussian curvature, *K*(*θ*, *ρ*), against the azimuthal angle, *θ*, at specific geodesic distances along the curved surfaces, *ρ*, show that for domes the Gaussian curvature is on average positive (〈*K*〉 > 0) and grossly radially symmetric, except for some defects around the gel edge (Fig. 2C). For wrinkled shapes the average Gaussian curvature is negative (〈*K*〉 < 0), and *K*(*θ*, *ρ*) oscillates between positive and negative values with amplitudes larger than the average that decay to vanishing values as we retract from the sheet boundary. This plot also clearly demonstrates that the wrinkled shapes have a single undulation mode, with no apparent higher order forming buckles. Plotting the measured perimeter on the curved surface against the geodesic distance and comparing it to the perimeter of the projected (flat) surface, *βρ* (*β* = 2*π* for a perfect disc, Fig. S5), shows that for domes the perimeter increases slower than linearly while for wrinkled shapes the perimeter increases faster, as expected in view of the increase in the oscillation amplitudes as the sheet boundary is approached (Fig. 2D).

**Figure 2.**
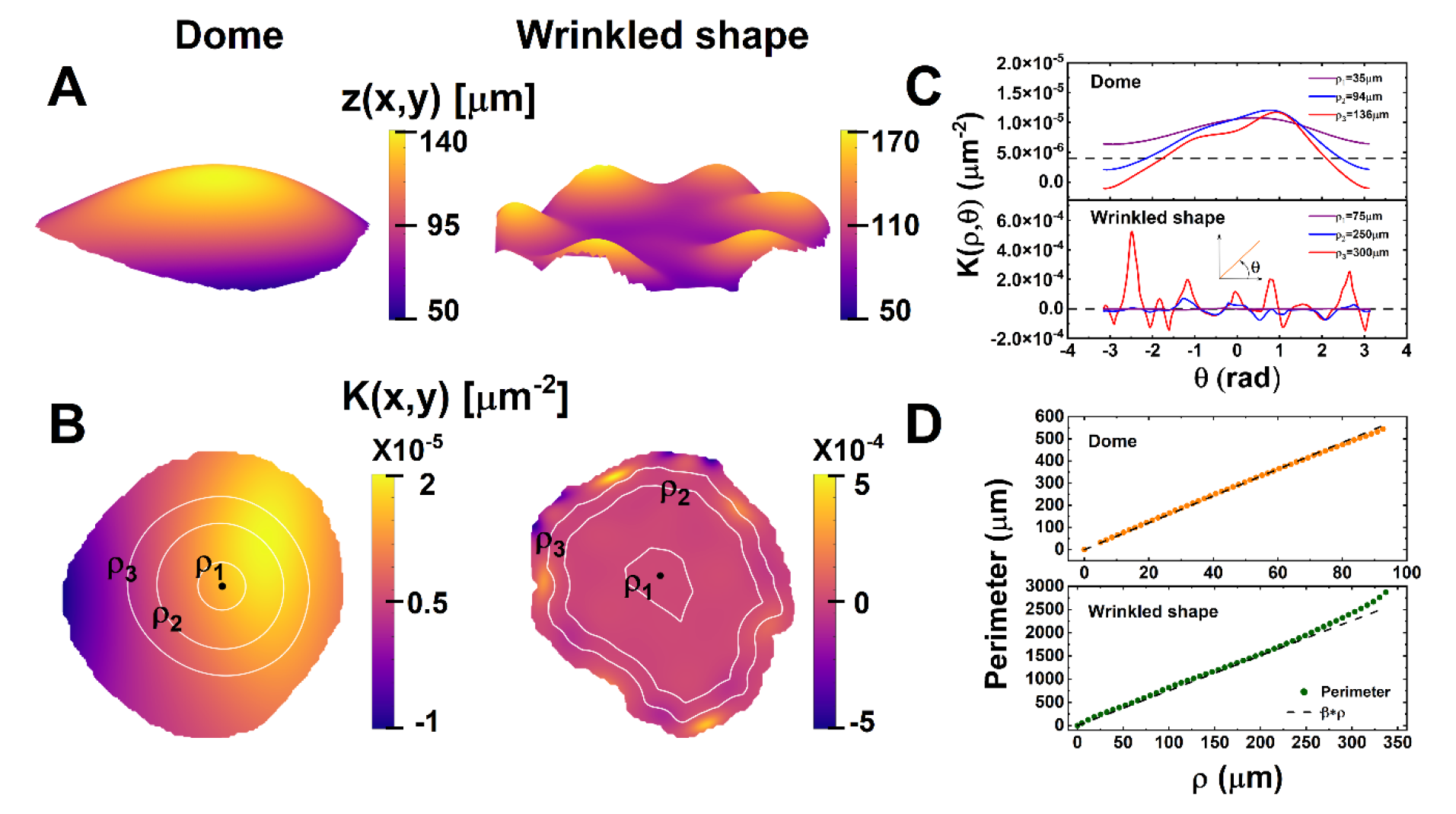
3D shapes geometry - domes *vs.* wrinkled shapes. **A)** Upper surface (smoothed) topography *z*(*x*, *y*) and **B)** top view 2D map of the measured local Gaussian curvature, *K*(*x*, *y*), for a dome (left) and a wrinkled shape with *m* = 6 nodes (right). Lines of constant geodesic distances, *ρ*, are highlighted in white. The black dot defines the center (*ρ* = 0) of the surface. **C)** The measured local Gaussian curvature along the three geodesic distances in (B) against the azimuthal angle, *θ*. Spatial average values of 〈*K*〉 (dashed line): 6 · 10^−6^ μm^−2^ (dome) and −6 · 10^−7^ μm^−2^(wrinkled shape). **D)** The measured (‘3D’) perimeter against the geodesic distance compared to the perimeter of the flat (‘2D’ projected) gel, *βρ* (dashed line). *β* ≅ 2*π* (dome) and *β* ≅ 1.2*·*2*π* (wrinkled shape).

### Shape selection is scale-free and encoded in the disc radius to thickness aspect ratio

We can now turn to ask how the final 3D configurations vary with the system initial aspect ratio? To achieve a two-order magnitude variation in initial aspect ratio, 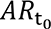 = 2 − 50, we fabricate gels in a range of initial thicknesses (65 − 330 µm) and radii (1200 − 6500 µm). The diverse shapes collapse on a universal linear line (slope ∼0.4, see also Fig. 1K), where domes, intermediate, and wrinkled shapes, segregate to low, intermediate, and large initial (and final) aspect ratios, respectively (Fig. 3A and B). For 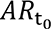 ≥ 10, the increase in aspect ratio produces a set of shape transformations, expressed in refinement of the wrinkled configurations (Fig. 3B). While different combinations of 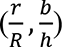 could produce the same net reduction in aspect ratio, we find that *all* systems, regardless of their final configuration and dynamic pathway, exhibit the same net radial (*r*/*R* ≅ 0.1) and vertical (*b*/ℎ ≅ 0.25) shrinking (Fig. 3C). This infers that all gels eventually compact similarly and thus undergo the same relative volume drop. This signifies that these are intrinsic material properties which do not depend on the system dimensions and geometry, but are determined by the actin, crosslinker, and myosin densities, which are the same for all the studied systems. Those systems thus having the same initial aspect ratio adopt the same steady state 3D configuration, independent of their actual initial dimensions, indicative of shaping scalability.

**Figure 3.**
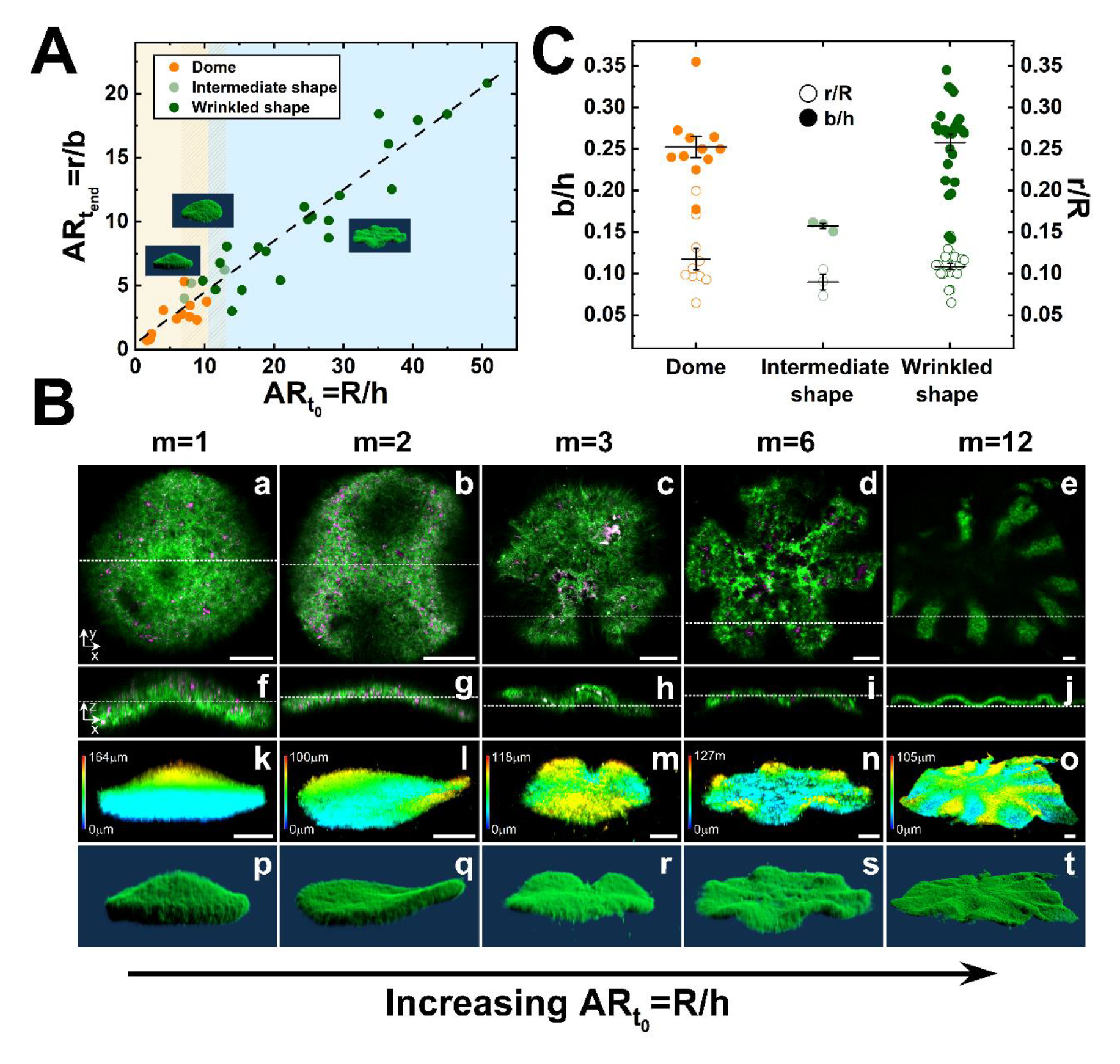
Shape selection is scale-free and encoded in the disc radius to thickness aspect ratio. **A)** 3D shapes phase space against the initial (and final) aspect ratios with representative images. **B)** Confocal microscopy images of final selected shapes against the initial aspect ratio. Top (a-e) and side views (f-j) confocal images and 3D representations, showing height maps (k-o) and surface plots (p-t), of a dome (a,f,k,p; 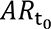 = 7.9), a 2-peak saddle (b,g,l,q; 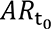 = 9.8), and wrinkled shapes with 3 (c,h,m,r; 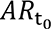 = 15.4), 6 (d,i,n,s; 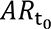 = 17.8), and 12 (e,j,o,t; 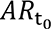 = 40.7) nodes. Top views in a,b,c,d, and e, are measured along the dotted lines in f,g,h,i, and j, respectively. Side views in f,g,h,i, and j are measured along the dotted lines in a,b,c,d, and e, respectively. Actin (green) and myosin (magenta). Scale bars: 100 μm. **C)** Net contraction in radial, 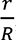, and vertical, 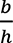, directions for the three main shapes. Shown are individual data points and their corresponding mean ± SD values.

### The final 3D configuration dependence on disc thickness and radius

We can now turn to ask what is the explicit dependence of the number of nodes *m* on the disc initial thickness and radius? This dependence can be estimated from the ratio of the measured steady state projected perimeter, *p* = *βr*, and the wrinkles wavelength *λ*. For uniaxially stretched thin elastic sheets of thickness *b*, the minimization of the stretching and bending elastic energies yields *λ*∼*L*^0.5^*b*^0.5^ (28, 29), where *L* is the length of the wrinkled domain perpendicular to the wrinkling direction. For discs, this ‘penetration’ length describes how far the undulations propagate radially into the sheet (Fig. 4A), and it can be evaluated from the deviation of the local Gaussian curvature from its average value squared, Δ*K*^2^(*r*, *θ*) ≡ (*K*(*r*, *θ*) − 〈*K*〉)^2^, which oscillates azimuthally matching the sheet wrinkling deformation. The measured (mean) penetration length *L* thus corresponds to the distance from the boundary where Δ*K*^2^(*r*) ≡ 〈(*K*(*r*, *θ*) − 〈*K*〉)^2^〉_*θ*_, the azimuthal average of Δ*K*^2^(*r*, *θ*), vanishes (Fig. S6). In contrast to uniaxially stretched sheets (28), both *L* and *λ* are insensitive to system size, i.e., final projected radius *r* (Fig. S7). This may indicate that radial and/or azimuthal tensile stresses develop in the sheet, confining the wrinkles to the sheet boundary. We find thus that *L* and *λ* are set merely by the sheet thickness (Fig. 4A). Explicitly, *L*∼*b* (inset, Fig. 4A), and *λ* = *αb*^0.5^ indicating that thinner sheets are more prompt to bend. *α* is a numerical pre-factor, independent of *r* (Fig. S7B). Interestingly, the same scaling relation for *λ* has been found for pre-programmed, externally stimulated, thin elastic chemically crosslinked gel discs (30).

**Figure 4.**
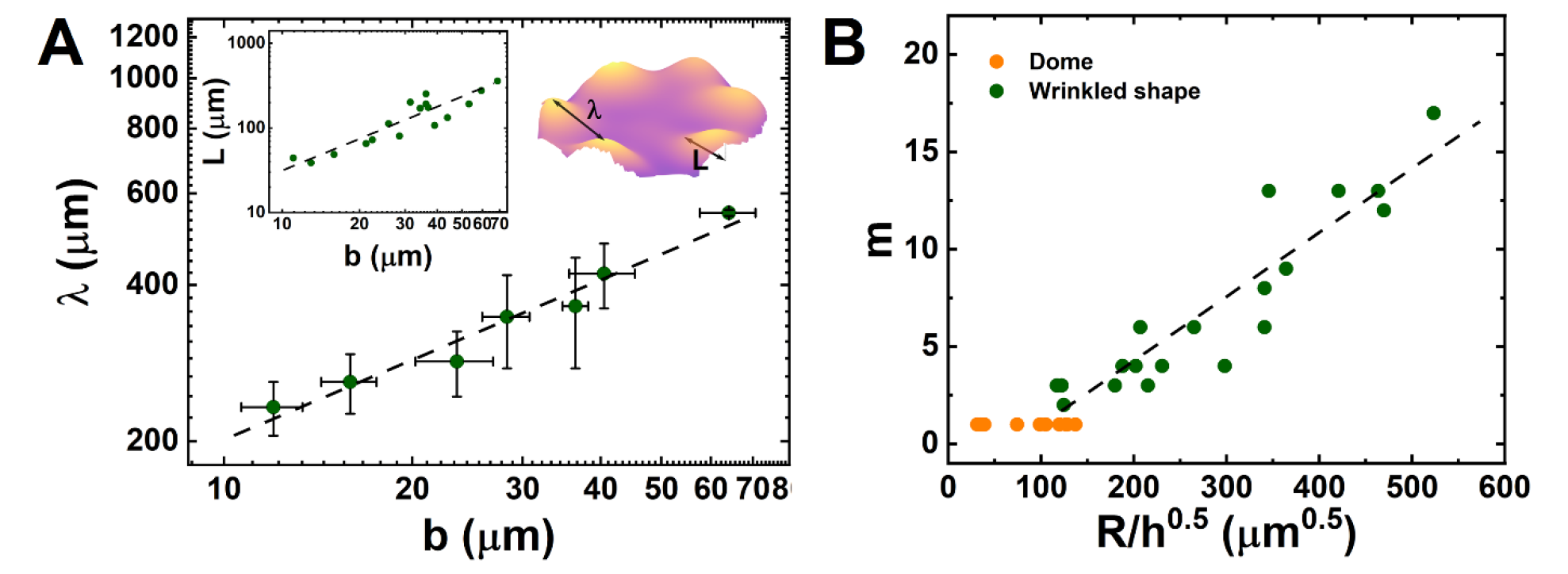
Wrinkled shapes final configuration. **(A)** Log-log plots of the measured wrinkles wavelength (mean ± SD, *N* = 3 − 4 gels with variable final radii) and penetration length (inset) against the sheet thickness yield *λ* = *αb*^0.51±0.04^ with *α* = 64 *μm*^0.5^ and *L*∼*b*^1.2±0.14^. Image: *L* and *λ* are illustrated. **(B)** The number of wrinkles grows linearly with 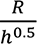 with a slope of 0.033 ± 0.003 (mean ± SD). Data for domes (*m* = 1) is also depicted.

Altogether, these relations yield: 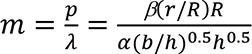, suggesting that 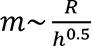, in accord with our experiments (Fig. 4B, green dots). The pre-factor, 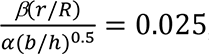, estimated from the parameter values, 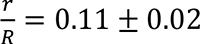 and 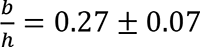 (Fig. 3C), and *β* = 2*π*(1.2 ± 0.02) (mean ± SD), averaged over all gels forming wrinkled shapes, agrees very well with the one extracted from the fit (dashed line). While this scaling is derived for wrinkled shapes, domes fall on the continuation of that same line, signifying that the intrinsic active stresses and the gel material properties governing shape deformation, are determined by the actin, crosslinker, and myosin densities, the same for all the studied systems.

## Discussion

Here, we employ an *in vitro* approach to determine the selection rules that govern shape deformation driven by contractile stresses in living systems. This approach presents a huge advantage as it allows us to reveal these selection rules under well-defined and controlled conditions, without the complexity of living systems. We access these selections rules by using a set of intrinsically identical, active (not-prepatterned) elastic actomyosin gel discs of variable initial dimensions and geometry, recreated using the same actin and myosin building blocks of the cell cytoskeleton. Using this system, we demonstrate that a family of 3D shapes can arise with the same set of molecular building blocks and that shape selection is encoded in system geometry and not on its actual dimensions. These evidence, may further indicate a universal, scale-free, shape selection mechanism, underlying the shape of cells as well as multi-cellular contractile tissues.

Our results suggest that while the dynamic pathways may depend on the poroelastic fluid dynamics (8, 18) and on the detailed interactions of the different microscopic components, the final shapes seem to be predicted by the general theory of elastic deformations of thin sheets (27). However, unlike passive elastic systems where the favored shape is always the minimal energy configuration for a specified distribution of forces, here the distribution of stresses can change as the system continues to deform until it reaches a mechanically stable state (29).

It remains to be explored how the self-organized structural order of actin filaments within the gel set up the gradient in active stresses (azimuthal *vs*. radial) that drive shape deformation. Moreover, how the dynamic pathways of network contraction, distinct for gels ending up in domes and wrinkled shapes, depend on the poroelastic fluid dynamics.

From practical purposes, these bio-inspired active materials present huge advantage towards the development of bio-compatible elastic active materials with controllable, molecularly induced, active stresses. These design principles can then be used to create desired, soft robots of target shapes, that are both tunable and robust.

Altogether, these results provide novel perspectives on the mechanically induced spontaneous shape transitions in contractile active matter and can help to uncover new mechanisms that drive shape selections in living systems across scales. Furthermore, it can help to understand how shape selections in living systems associate with their functions.

## Materials and Methods

### Protein purification

G-actin is purified from rabbit skeletal muscle acetone powder by gel filtration (31), stored on ice, and used within 2-3 weeks. Actin is labeled on Cys374 with Alexa-Fluor 488. Fascin is produced as a GST-fusion protein. Myosin II skeletal muscle is purified following (32) and labeled with Alexa-Fluor 647 at pairs of engineered cysteine residues (14, 33).

### Preparation of intrinsically contractile elastic actomyosin gel discs

Actomyosin gel discs are prepared by polymerizing 5 μM G-actin with 16.7 nM myosin II (in large aggregates of ∼150 myosin dimers), 280 nM fascin, and 2mM ATP (see also (18, 25) and SI appendix). We use an ATP-regenerating system and an anti-bleaching solution. The percentage of labeled G-actin and myosin II is 5 mol%. A drop of that solution is squeezed between two PEG-passivated coverslips, placed in a homemade sample holder (25). We generate disc of variable height ℎ and radii R, by varying the drop volume and spacing between the two coverslips.

### Microscopy techniques

Samples are imaged using a laser scanning confocal microscope (LSM 880, Zeiss Germany) in fast Airyscan mode using a ZEN software (Zeiss Germany). We minimize the acquisition time between frames by working in single channel mode. The 3D steady state shapes are imaged in standard Airyscan mode to optimize sampling resolution. Actin is excited at 488 nm and myosin at 647nm.

### Data quantification

From the confocal images we extract for each time point: the disc radius *r*(*t*) and thickness, *e*(*t*), from which the variation in disc aspect ratio, *AR*(*t*) = *r*(*t*)/*e*(*t*), and radial and vertical strains, 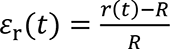 and 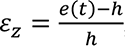 are derived. Also extracted is the local filament orientation. The results are presented in color map in HSB mode (34). Additional measured parameter values include: the spacing between the two coverslips, *h*, the disc initial radius, *R*; the system final projected radius *r* and thickness *b*; for wrinkled shapes: the wrinkles number, *m*, wavelength, *λ*, and penetration depth, *L*. Data is corrected for index refraction mismatch between the objective immersion liquid and the aqueous sample (35).

#### 3D shapes geometry

For each final shape, the topography *z*(*x*, *y*) of the top surface is generated from a stack of *xy* cross-section confocal images. We match a semi-geodesic coordinate system (*ρ*, *θ*) to each such surface. On that surface we calculate the local Gaussian curvature, *K*(*θ*, *ρ*), using the first and second fundamental forms (36), extract its spatial average value 〈*K*〉. We calculate the (‘3D’) perimeter *p*(*ρ*) at each geodesic distance *ρ*, which we calculate using the Heat diffusion method (37). Similarly, the projected ‘2D’ perimeter *p*(*r*) = *βr* is extracted by matching the surface to a polar coordinate system (*r*, *θ*). The local projected radial distances *r* are computed using the two-dimensional Multistencil Fast Marching Method (MSFM2D) (38). The parameter, *β*, measures the surface non-circularity (*β* = 2*π* for a perfect disc). The wrinkles penetration depth *L* is evaluated from the azimuthal average of the standard deviation of the local Gaussian curvature Δ*K*^2^(*r*) ≡ 〈(*K*(*r*, *θ*) − 〈*K*〉)^2^〉_*θ*_.

Data quantification is performed using MATLAB (MathWorks, MA, USA), Zen Black 2.1 (Zeiss, Germany), Huygens Professional software package (Huygens; Scientific Volume Imaging, Hilversum, Netherlands), Origin (OriginLab Corp., MA, USA), and ImageJ software.

Due to space limitation a comprehensive Method section is provided in the SI Appendix.

## Acknowledgments

We thank Sam Safran, Haim Diamant, Ido Levin, and Eran Sharon for useful discussions. We thank Dina Aranovich for protein purification and labelling. G.L. is grateful to the Israel Ministry of Science and Technology for the Jabotinsky PhD Scholarship. A.B.G. is grateful to the Israel Science Foundation (grant 2101/20) and the Ministry of Science and Technology (grant 3-17491) for financial support.

## Contributions

L.G. Performed experiments, developed all analytical methods for data quantification, analyzed the experimental results, prepared the Figures and Movies, and wrote the manuscript.

G.S. Designed and performed experiments.

A.S. Developed analytical methods for data quantification.

A.B.G. Developed the experimental system and wrote the manuscript.

## Data availability

The data that support the findings of this study are available from the corresponding author upon reasonable request.

## Supplementary appendix

### Materials and Methods

#### Protein purification

G-actin is purified from rabbit skeletal muscle acetone powder by gel filtration (1) (HiPrep^TM^ 26/60 Sephacryl^TM^ S-300HR, GE Healthcare), stored on ice, and used within 2 weeks. Actin is labeled on Cys374 with Alexa-Fluor 488. Fascin is produced as a GST-fusion protein. Purification of myosin II skeletal muscle is performed following standard protocols (2). Myosin II is labeled with Alexa-Fluor 647 at pairs of engineered cysteine residues (3, 4).

### Experimental procedure

#### Preparation of intrinsically contractile elastic actomyosin gel discs

Macroscopically contractile elastic actomyosin networks are formed by polymerizing G-actin with myosin motors, added in the form of large aggregates (∼150 myosin dimers/aggregates), the strong crosslinker fascin, and ATP (see also (5, 6)). Specifically, the solution contains 10 mM HEPES pH = 7.0, 1 mM MgCl_2_, 25 mM KCl, 2 mM ATP, kept constant using an ATP-regenerating system (0.5 mg mL^−1^ creatine kinase and 5 mM creatine phosphate), 200 μM EGTA, an anti-bleaching solution (0.1 mg mL^−1^ glucose oxidase, 0.018 mg mL^−1^ catalase, and 5 mg mL^−1^ glucose), 5 μM G-actin, 280 nM fascin, and 16.7 nM of myosin II. The percentage of labeled G-actin and myosin II is 5% (molar percentage).

A drop of that solution is squeezed between two PEG-passivated coverslips, placed in a homemade sample holder which fits the dimensions of a standard microscope stage (6). We use Teflon, Parafilm, or tape spacers to control the spacing between the two coverslips, through which the drop height ℎ, which defines the gel initial thickness, is controlled. By varying the drop volume and spacer type we generate discs of controllable initial radii *R*. In our experiments, the gel initial thickness and radius range between 65 and 330 µm and 1200 and 6500 µm, respectively.

### Microscopy techniques

Samples are imaged using a laser scanning confocal microscope (LSM 880, Zeiss Germany) in fast Airyscan mode using a ZEN software (Zeiss Germany). We minimize the acquisition time between frames by working in single channel mode (i.e., the actin channel, Ex. wavelength of 488 nm). Confocal images of the 3D steady state shapes are collected in standard Airyscan mode to optimize sampling resolution, imaging both actin (Ex. 488 nm) and myosin (Ex. 647nm(.

### Data quantification

#### Analysis of the contraction and buckling dynamics of the gel discs

Laser scanning confocal microscopy imaging is used to follow the contraction and buckling dynamics of the gels with time and image their 3D shape at the end of the contraction process when the gels reach a mechanical stable (steady) state. To quantify the contraction dynamics, the temporal radius *r*(*t*) and (mean) temporal thickness, *e*(*t*), are extracted from the confocal images. Then, the variation in gel aspect ratio, *AR*(*t*) = 2*r*(*t*)/*e*(*t*), and radial and vertical strains, 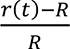 and 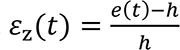 are derived at each time point. The distribution of actin density normal to the thickness is also extracted from the confocal images as well as the in-plane local actin filaments alignment in the network structure. Finally, we extract parameters value including the spacing between the two coverslips, *h*, which sets the gel initial thickness, the drop (i.e., gel) initial radius, *R*, the final projected radius *r* and thickness *b*, the number of wrinkles per gel, *m*, and the mean wrinkles’ wavelength, *λ*. Data quantification is performed using MATLAB (MathWorks, MA, USA), Zen Black 2.1 (Zeiss, Germany), Huygens Professional software package (Huygens; Scientific Volume Imaging, Hilversum, Netherlands), Origin (OriginLab Corp., MA, USA), and ImageJ software.

##### Temporal projected radius r(t)

From practical purposes, i.e., accessible field of view, quantification of the contraction and buckling dynamics are restricted to gels with initial diameters 2*R* ≤ 3800 µm, where *R* is the initial drop radius. We extract the temporal projected radius *r*(*t*) from confocal, *xy* cross-section, images by fitting the gel area to a circle. If less than a quarter of the gel is visible in the field of view, which usually happens at the very initial stages of gel contraction, the radius is evaluated from the changes in the position of the gel boundary between two sequential time points. For gels forming a dome shape, this analysis is performed in maximal intensity projection (MIP) mode, such that the gel widest projected diameter is probed at each instant.

##### Temporal mean thickness e(t) and final thickness b

We evaluate the temporal mean thickness *e*(*t*) from, *xz* and *yz* cross-sections, confocal images and use it to calculate the temporal vertical strain *ε*_z_(*t*). For each time point, we use between 25-50 data points, localized at different positions along the gel surface, where each value corresponds to the normal distance between the top and bottom, surfaces at that position. Errors correspond to standard deviations of the experimental values. For gels forming wrinkled and intermediate shapes these data points span the whole gel sheet. For gels ending up in a dome shape, thickness measurement is restricted to the central region of the disc up to the time point the thickness becomes homogeneous everywhere. The thickness is extracted from confocal microscopy images in standard imaging mode except for gels forming wrinkled and intermediate shapes. For these gels, the thickness is extracted from maximal intensity projection (MIP) images, which allows to resolve the gels’, top and bottom, surfaces during vertical contraction. Once the gel becomes sufficiently dense, which usually corresponds to the onset of the lateral contraction phase, the thickness is measured in standard imaging mode. The final steady state thickness *b* is evaluated similarly on images acquired after the gels have reached a mechanically stable state (*t*_*end*_). Data is corrected for index refraction mismatch between the objective immersion liquid and our aqueous sample by multiplying the measured thickness by 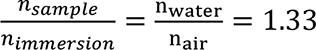 (7).

##### Drop initial thickness ℎ and radius R

These parameter values are measured on images acquired after the systems have reached a mechanically stable state (*t*_*end*_). The gel initial thickness ℎ corresponds to the distance between the two coverslips, determined from side-view, *xz* or *yz* cross-section, confocal images. Data is corrected for index refraction mismatch between the objective immersion liquid (here air) and the aqueous sample 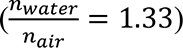 (7). To evaluate the gel (drop) initial diameter, 2*R*, multiple top-view, *xy* cross-section, confocal images are acquired with 10% overlap and stitched into a single image. The number of images used for stitching (usually 7×7) depends on the size of the drop and objective magnification. The drop (and gel) initial radius *R* is determined by fitting the drop perimeter to a circle.

##### Density distribution of actin normal to the thickness

For a given time point, the average density distribution of actin, normal to the gel, top and bottom, surfaces is calculated from 15-25 individual density profiles, each 30-pixel in width. Errors correspond to standard deviations of the experimental values.

##### In plane local orientation of the actin filaments/bundles

For a given time point, the in plane local orientation of the filaments is evaluated using the OrientationJ Analysis plugin in ImageJ via which a structure tensor, characterizing the filaments’ mean orientation and isotropy in a local window, is computed. The local window is characterized by a 2D Gaussian function of standard deviation σ, of the size of the structure of interest (e.g., thickness of the filament/bundles). The results are presented in color map in HSB mode, where hue corresponds to mean local orientation, saturation is coherency, and brightness is the source (original) image (8).

##### 3D steady state shapes: Extracting the topography z(x, y) and the local Gaussian curvatures K(x, y)

This analysis is performed on gels having reached a mechanically stable state (*t*_*end*_). For each gel, the topography *z*(*x*, *y*) of the top surface is reconstructed from a stack of, initially binarized, *xy* cross-section confocal images. This surface, which has a unit-pixel depth and a resolution of 1 μm in the lateral *x*, *y* plane, is smoothed by fitting it to a 2D polynomic equation using ‘polyfit2D’ function in MATLAB (MathWorks, MA, USA) using (*N* + 1) · (*N* + 2)/2 polynomial coefficients(9). We find empirically that for all gels the smoothed surface optimally fits the actual gel top surface for *N* = *m* + 2, where *m* is the number of nodes of an actual gel. This smoothed surface is assumed to reliably represent the shape and local curvatures of the gel sheet mid-surface at steady state. This is explicitly true for wrinkled and intermediate shapes which form thin sheets (i.e., *b*/*r* ≪ 1). For domes, which are relatively thicker (i.e., *b*/*r* < 8), these values provide a lower bound estimate of the local and average, Gaussian curvatures. The local Gaussian (*K*(*x*, *y*)) curvature is then computed at each point along that smoothed surface, using the second and first fundamental forms II and I(10).

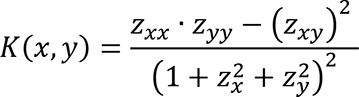

The average Gaussian curvature 〈*K*〉 is the number average of the local Gaussian curvature values.

##### Distribution of the local Gaussian curvature as a function of the azimuthal angle for different geodesic distances

To compute the distribution of the local Gaussian curvature, *K*(*θ*, *ρ*), against the azimuthal angle, *θ*, at specific geodesic distances along the curved surfaces, *ρ*, we match a semi-geodesic coordinate system (*ρ*, *θ*) to the polynomial-fitted smoothed surfaces. We compute the local geodesic distances on the deformed surfaces by using the Heat diffusion method (11) with an optimal time step of *t* = 10. For dome shapes, the local geodesic distance is calculated relative to the gel center *ρ*(*x*, *y*). The gel center is thus defined as *ρ* ≡ 0. In that case, the gel center is defined at the location of the highest point (in z-axis) of the smoothed surface. In wrinkled shapes the buckling instability occurs at the gel periphery; the undulation amplitude decays as we retract from the gel periphery. Thus, for these gels it is more natural to calculate the geodesic distances from the boundary, *ρ*^′^(*x*, *y*). The gel boundary (edge) is detected using ‘bwboundaries’ (MATLAB library function) and is defined as *ρ*^′^ ≡ 0. The sheet center now corresponds to the location of the maximal geodesic distance 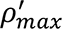, from which the local geodesic distance measured relative to the sheet center can be evaluated, 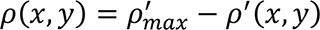. Note, that choosing the gel boundary as the origin for calculating the geodesic distances allows to evaluate the sheets perimeter (projected and actual) and quantify its deviation from perfect circularity (disc). In our experiments, the deviation from perfect circles is usually observed for gels with initial diameters larger than 4200 μm which are more prompt to deform upon squeezing.

##### Non-circularity coefficient β

For each gel we extract the parameter, *β*, which provides a measure of the non-circularity of the gels. This parameter is evaluated from the slope of the linear fit of the function relating the measured projected (2D) perimeter *p* and the projected radial distance, *r*, that means *p* = *βr*. Note that for a perfect disc *β* = 2*π*. The local projected radial distances are computed using the two-dimensional Multistencil Fast Marching Method (MSFM2D) (12). Here we employ a similar strategy as done for calculating the geodesic distance. That is, for dome shapes the local projected radial distance is calculated relative to the gel center, *r*(*x*, *y*). For wrinkled shapes, the local projected radial distance is first calculated relative to the sheet boundary *r*′(*x*, *y*). Then, the local projected radial distance measured from the gel center is evaluated at each point from 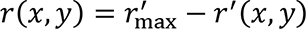 where 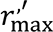 is the maximal projected radius calculated from the sheet boundary. For each gel, the changes in measured projected (2D) perimeter, *p*(*r*), is evaluated by creating contour lines along the surface at various projected radial distances *r*, using the ‘contour’ MATLAB library function, and computing the length of each contour using ‘contourlengths’ (MATLAB library function).

##### Measured perimeter profile vs. the geodesic distance ρ

The measured perimeters on the deformed surface (so-called 3D perimeters) at different geodesic distances, *p*(*ρ*), are extracted similarly. This is done by creating 3D curves representing all the topographic points *z*(*ρ*) at a given geodesic distance *ρ*, and computing the length of this contour line using the ‘arclength’ MATLAB function(13), which compute the arc length of a space curve represented as a sequence of points. These contour lines are created using the ‘contour’ MATLAB library function and then, the topographic points along these contours are extracted by ‘C2xyz’ MATLAB function (14).

##### Wrinkles penetration depth L

We use the square of the deviation of the local Gaussian curvature from its average value, Δ*K*^2^(*r*, *θ*) ≡ (*K*(*r*, *θ*) − 〈*K*〉)^2^, to evaluate the wrinkles penetration depth, *L*. *θ* is the polar angle and 〈*K*〉 is the average measured Gaussian curvature. These local values are then averaged along the polar angle *θ* for each projected radial distance *r*, yielding Δ*K*^2^(*r*) ≡ 〈(*K*(*r*, *θ*) − 〈*K*〉)^2^〉_*θ*_. The wrinkles penetration depth *L*, defines the distance from the boundary at which Δ*K*^2^(*r*) vanishes and it corresponds to the local minima of Δ*K*^2^(*r*) *vs. r* in log-log representation. It provides an average value of the penetration depth of the wrinkles formed on a given sheet.

## Supplementary Figures

**Figure S1.**
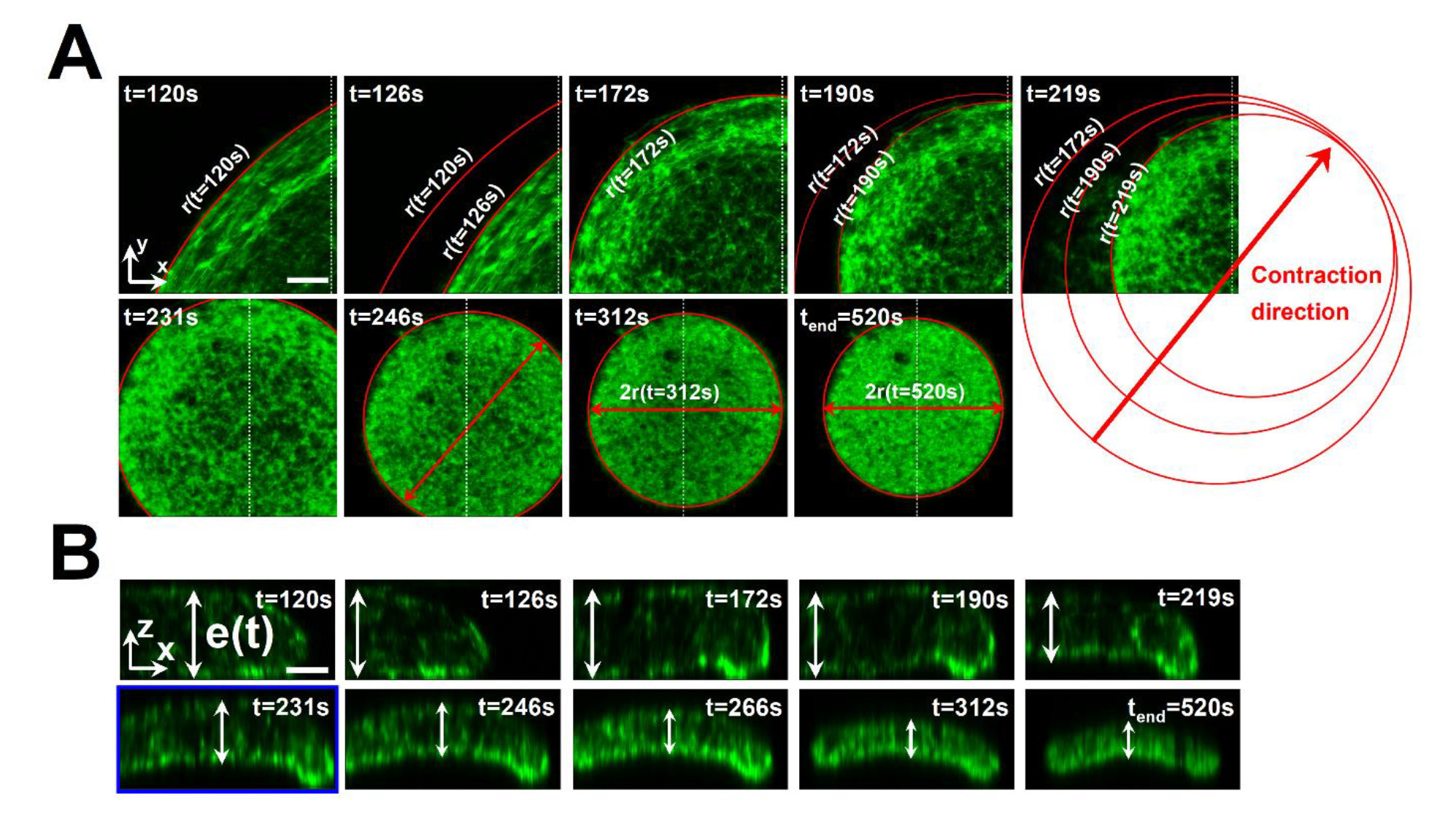
**Analysis of the contraction dynamics of the gel in Fig. 1C-E that self-organize into a dome shape. A) Extracting the temporal projected radius *r*(*t*)**. Shown are Maximal Intensity Projection (MIP) of top view, *xy* cross-section, laser scanning confocal images acquired in fast Airyscan mode of the gel at different time points. The temporal projected radius, *r*(*t*) is evaluated by fitting the gel area to a circle (red). If less than a quarter of the gel is visible in the field of view, the radius is evaluated from the changes in the position of the gel boundary between two sequential time points. **B) Temporal thickness e(t) and buckling onset.** Shown are side view *yz* cross-section confocal images (standard mode) measured along the white dotted lines in (A). The thickness *e*(*t*) is measured perpendicular to the gel, top and bottom, surfaces. For each time point, between 25-50 data points, localized at different positions along the gel surface, are used. Each value corresponds to the normal distance between the top and bottom, surfaces at that position (see e.g., white arrows). Errors correspond to standard deviations of the experimental values. The images are corrected for index refraction mismatch between the objective immersion liquid (here air) and the aqueous sample (see Methods). Thickness measurement is restricted to the central region of the disc up to the time point the thickness becomes homogeneous everywhere (*t* = 266 sec). The buckling onset (blue rectangle) corresponds to the time point the gel center adopts a non-vanishing curvature. The steady state thickness *b* is evaluated similarly, but on high resolution images acquired in standard Airyscan mode (not shown). Scale bars are 100 μm. Gel initial dimensions: ℎ = 245 μm and 2*R* = 3400 μm.

**Figure S2.**
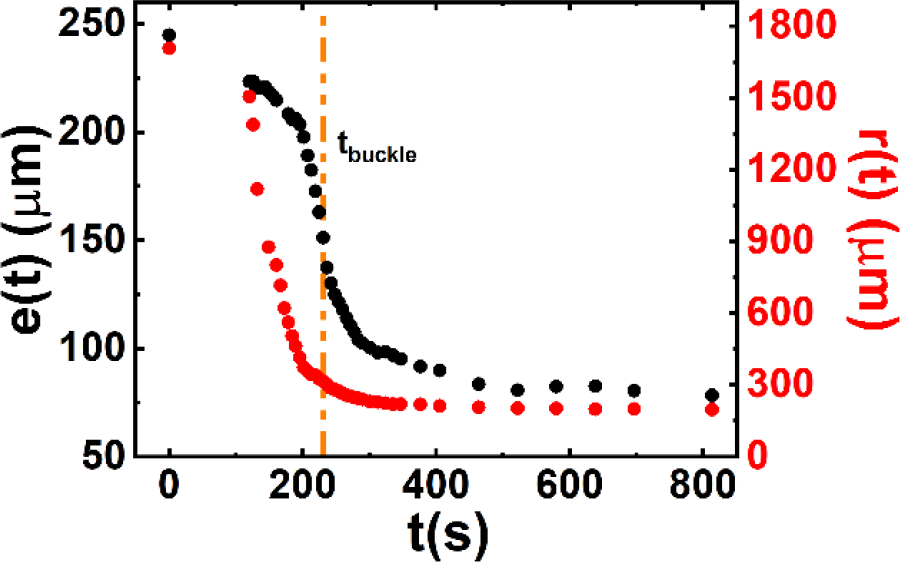
Temporal thickness *e*(*t*) and projected radius *r*(*t*) of the gel in Fig. 1C,D (main text) and Fig. S1. The orange dashed line marks the dome buckling onset (*t*_buckle_ = 231 sec). Gel initial dimensions: ℎ = 245 μm and 2*R* = 3400 μm.

**Figure S3.**
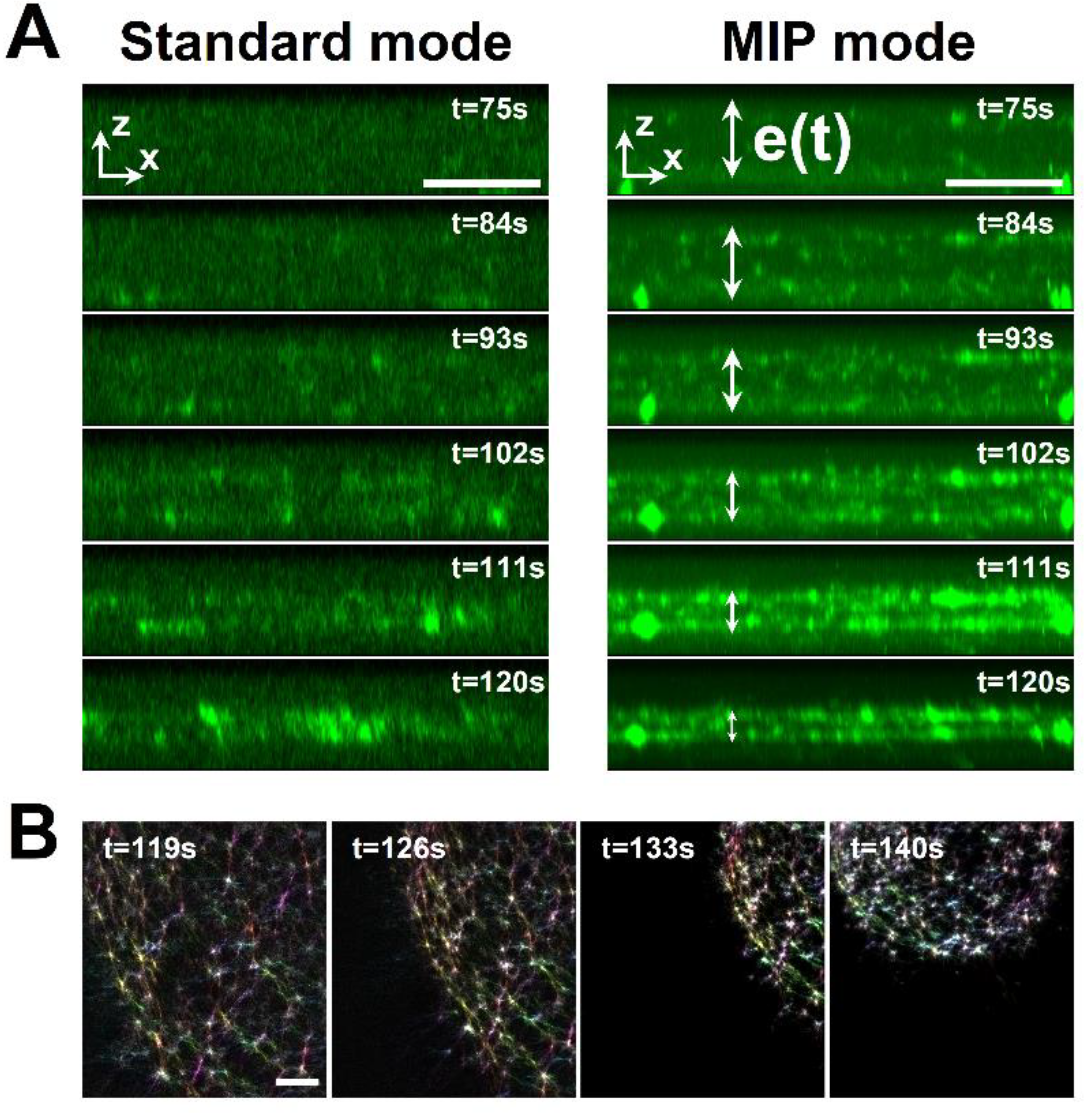
**(A) Thin sheet formation - probing the changes in temporal thickness for gels ending up in wrinkled and intermediate shapes.** Shown are laser scanning confocal side view images of the gel in Fig. 1F-H (main text) during the vertical contraction phase. The temporal thickness *e*(*t*) is extracted from Maximal Intensity Projection (MIP), *xz* and *yz* side view cross-section images (right column) as the thickness cannot be resolved in standard imaging mode (left column). *t* = 120 sec, marks the end of the vertical contraction phase and the initiation of the lateral (in-plane) contraction phase. At this point, the gel usually becomes sufficiently dense to be visualized in standard imaging mode. Gel initial dimensions: ℎ = 63 μm, 2*R* = 3260 μm (*m* = 4). **(B)** Top view, *xy* cross-section, laser scanning confocal images acquired in fast Airyscan mode of a gel ending in a wrinkled shape with 3 nodes, with the local filaments orientation, color coded to the orientational angle. The sheets remain structurally isotropic throughout radial contraction and wrinkling except for vanishingly small, aligned region of the actin bundles at the gel periphery. Gel initial dimension: ℎ = 80 μm, 2*R* = 3780 μm (*m* = 3). Scale bars in (A) and (B) are 100 μm.

**Figure S4.**
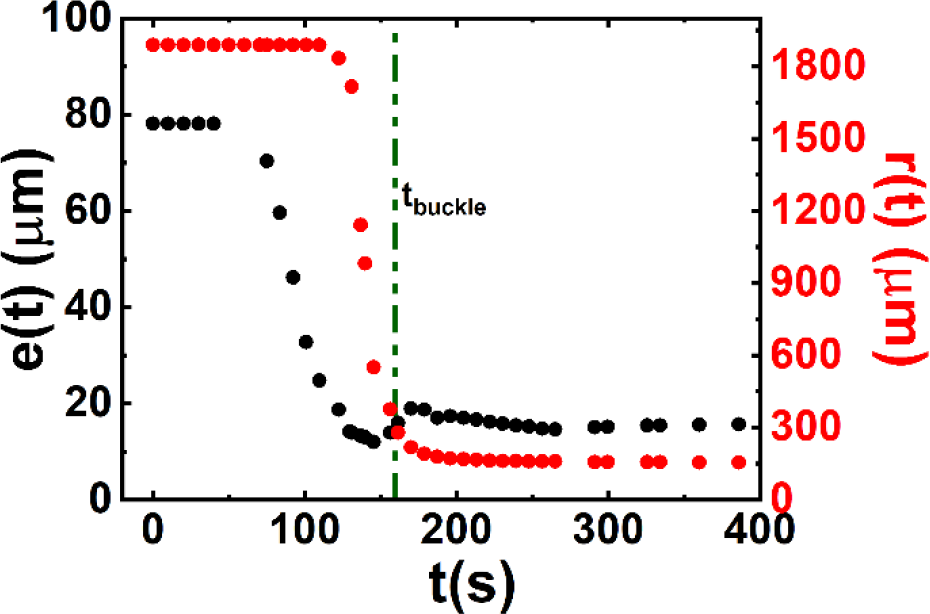
Temporal thickness *e*(*t*) and projected radius *r*(*t*) of the gel in Fig. 1G,H (main text). The green dashed line marks the wrinkling onset (*t*_buckle_ = 165 sec). Gel initial dimensions: ℎ = 63 μm, 2*R* = 3260 μm (*m* = 4).

**Figure S5.**
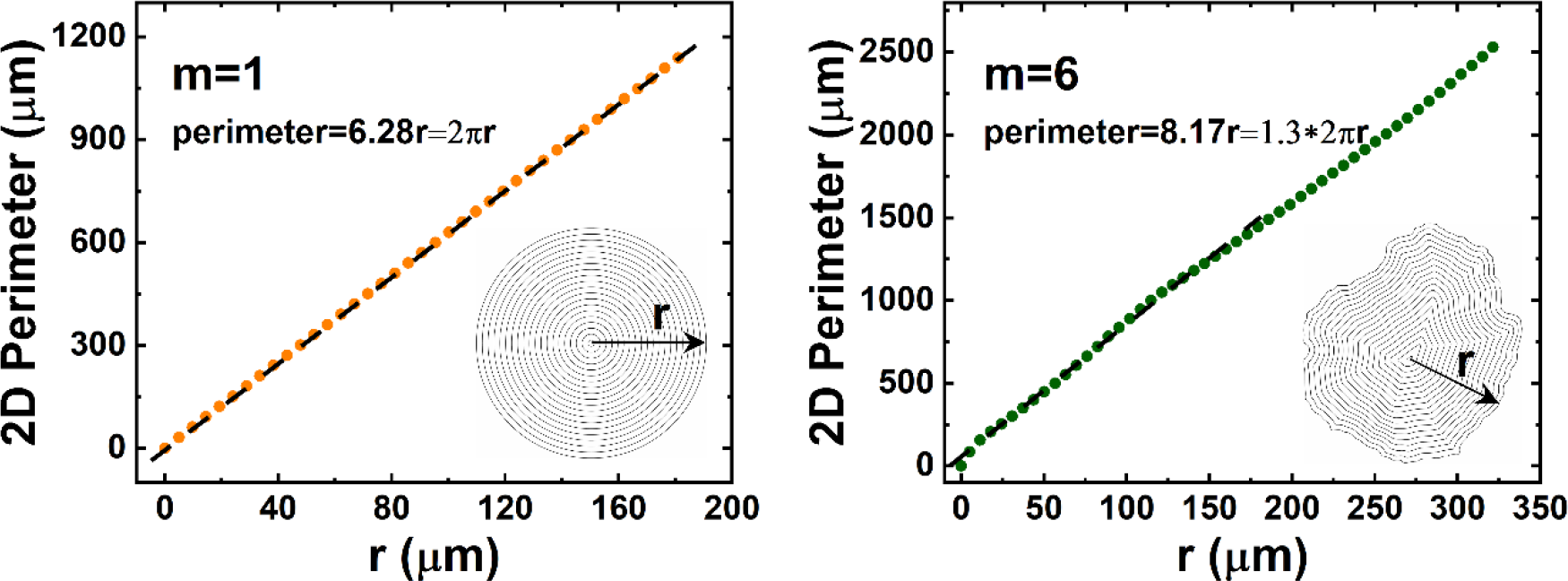
**Extracting the non-circularity coefficient *β* from projected perimeter**. Shown are the projected (2D) perimeter *p* against the projected distance *r* for the dome (left) and wrinkled shape (right) shown in Fig. 2. For a perfect circular disc, this relation *p* = *βr*, yields *β* = 2*π*. For each gel *β* is extracted by fitting the measured perimeter to a linear function. Inset: image of the gels’ contour lines of the projected perimeters at different projected distances *r*.

**Figure S6.**
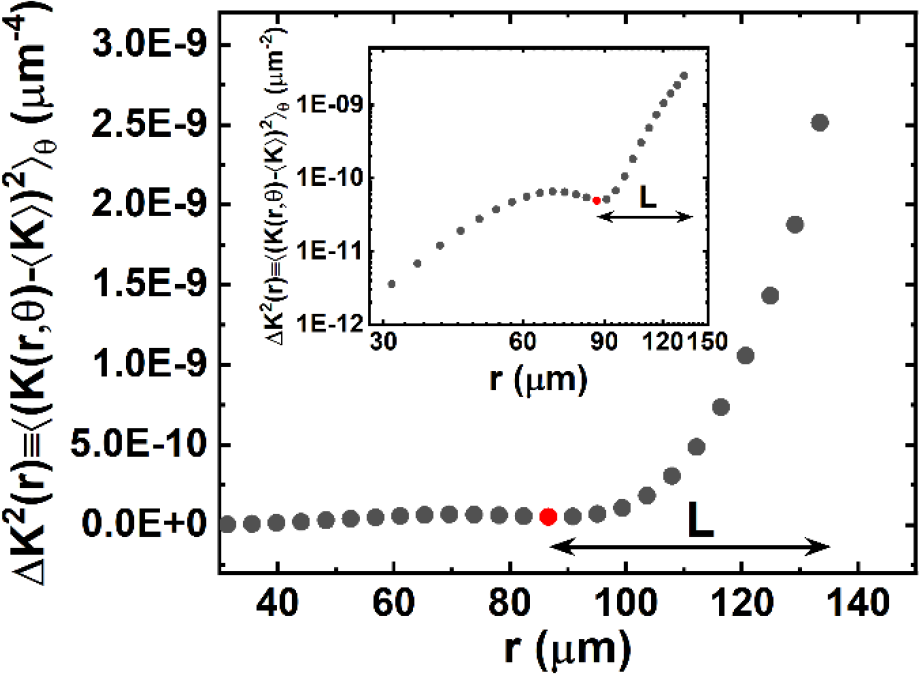
**Extracting the wrinkles penetration length *L***. Angular average of the deviation of the local Gaussian curvature from its average value squared, Δ*K*^2^(*r*) ≡ 〈(*K*(*r*, *θ*) − 〈*K*〉)^2^〉_*θ*_, against the projected radial distance *r* of the wrinkled shape in Fig. 2. The wrinkles penetration depth *L* defines the distance from the boundary at which Δ*K*^2^(*r*) vanishes (red dot). In log-log representation this corresponds to the position of the local minima (inset, red dot).

**Figure S7.**
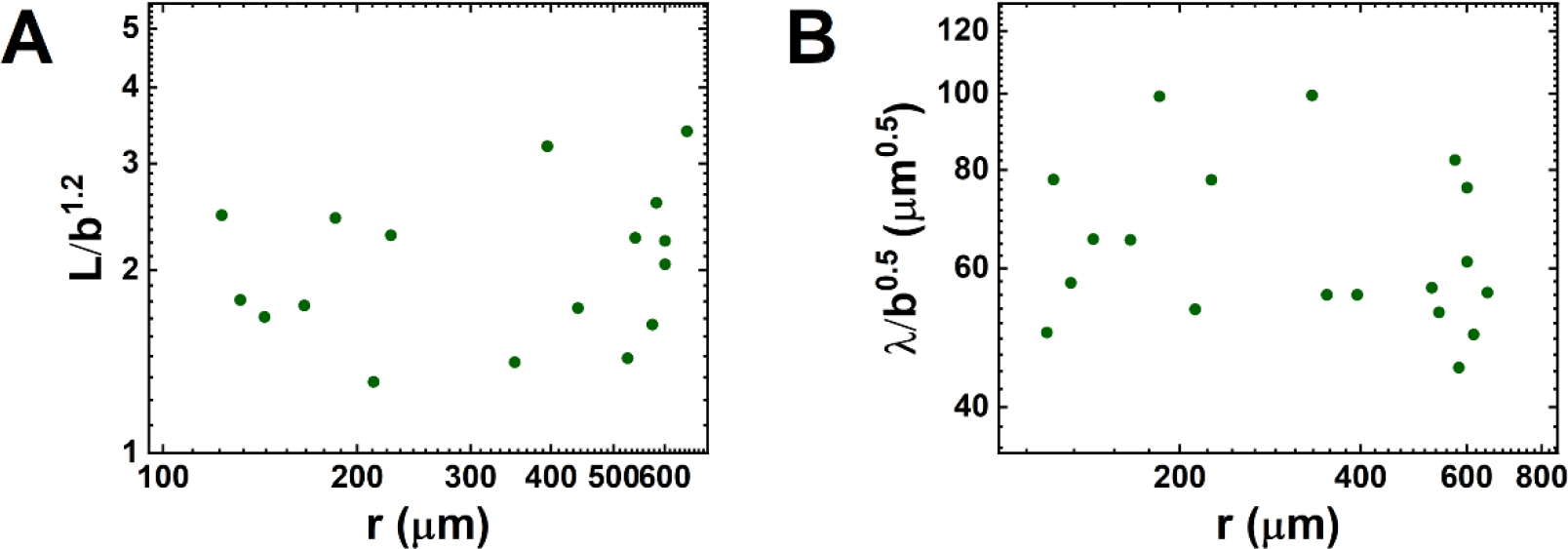
**Wrinkles penetration depth *L* and wavelength are independent of system size, i.e., steady state projected radius *r*. (A)** Measured wrinkles penetration depth *L* normalized by *b*^1.2^, which eliminates the dependence of *L* on the steady state thickness *b* (see Fig. 4A inset), for the various analyzed gels shows that the penetration depth is independent of system size (i.e., *r*). **(B)** Similarly, we find that the measured wrinkles wavelength *λ*, normalized by *b*^0.5^, is independent of the final project radius.

## Supplementary Movies

**Movie S1.** Contraction dynamics and buckling of a gel that self-organizes into a dome shape. Laser scanning confocal microscopy imaging of the gel shown in Fig. 1C,D (main text) and Fig. S1. Three views of the gel are shown: top view, *xy* cross-sections, measured along the dotted lines in the side view, *xz* (top) and *yz* (right) cross-section images. Similarly, side view, *xz* (top) and *yz* (right) cross-section images, are measured along the dotted lines in the *xy* cross-sections. Actin is labeled with Alexa-Fluor 488. Scale bar is 100 μm.

**Movie S2.** Local actin bundles orientation in the gel shown in Supplementary Movie S1. The results are presented in color map in HSB mode, where hue corresponds to mean local orientation, saturation is coherency, and brightness corresponds to the source (original) image. Scale bar is 100 μm.

**Movie S3.** Vertical contraction dynamics of the gel in Fig. 1F, H (main text) and Fig. S3A that self-organizes in a wrinkled shape. Shown are top view (*xy* cross-sections) and side view (*xz* (top) and *yz* (right) cross-sections) Maximal Intensity Projection (MIP) images of the gel. Actin is labeled with Alexa-Fluor 488. Scale bar is 100 μm.

**Movie S4.** Local actin filaments/bundles orientation of the gel in Fig. S3B that self-organize in a wrinkled shape with *m* = 3 nodes. The results are presented in color map in HSB mode, where hue corresponds to mean local orientation, saturation is coherency, and brightness is the source (original) image. Scale bar is 100 μm.

